# Computer-guided design of Z domain peptides with improved inhibition of VEGF

**DOI:** 10.1101/2024.11.29.626075

**Authors:** Carsten Geist, Abibe Useini, Aleksandr Kazimir, Richy Kümpfel, Jens Meiler, Christina Lamers, Stefan Kalkhof, Georg Künze

## Abstract

Computational protein design is becoming increasingly helpful in the development of new protein therapeutics with enhanced efficacy, specificity, and minimal side effects, for precise modulation of biological pathways. In vascular biology, the interaction between vascular endothelial growth factor A (VEGFA) and its receptors (VEGFR1–R3) is a pivotal process underlying blood vessel growth. Dysregulation of this pathway contributes to diseases such as cancer and diabetic retinopathy. Existing VEGFA inhibitors are effective but have limitations, driving interest in peptide-based therapeutics. Peptide inhibitors offer advantages, including reduced toxicity, improved formulation flexibility, and enhanced stability. This study leverages computational tools, particularly ProteinMPNN and Rosetta, to design optimized peptide-based VEGFA inhibitors. Building on the existing peptide templates mini-Z-1 and Z-1-2, new sequences were computationally predicted and experimentally validated. A novel peptide with improved affinity (K_D_ = 6.2 µM) compared to mini-Z-1 (K_D_ = 9.3 µM) was found, requiring only one round of design and testing. The integration of ProteinMPNN and Rosetta enabled a rapid and cost-effective pipeline for designing potent VEGFA inhibitors, underscoring the potential of computational peptide design in developing next-generation therapeutics targeting angiogenesis-dependent diseases.

## Introduction

Computer-aided drug design has become a powerful tool for developing compounds with improved therapeutic efficacy, selectivity, and reduced side effects (1). Moreover, computational methods not only accelerate the discovery of new lead compounds but also reduce the costs and time associated with traditional drug development. Precise drug treatment strategies have already been applied in several human disorders, including those involving dysregulated blood vessel growth (2).

Vascular endothelial growth factor A (VEGFA) plays a fundamental role in blood vessel formation during embryogenesis (vasculogenesis) and the growth of new vessels from existing ones (angiogenesis) (3,4). Vertebrate VEGFs are broadly expressed across tissues and are essential for blood vessel formation (5). They bind with high affinity to receptor tyrosine kinases (RTKs), specifically VEGF receptors 1–3 (VEGFR1-R3) (6). While VEGFR1–R3 share structural similarities, including seven extracellular immunoglobulin-like domains, a transmembrane helix, and a split tyrosine kinase domain, they differ in activation, signaling, and biological effects. VEGFs exist in various isoforms (VEGFA, B, C, D), with VEGFA being the most studied. VEGFA undergoes alternative splicing, influenced by factors like hypoxia and oncogenic signals, producing isoforms such as VEGFA111, VEGFA121, VEGFA165, VEGFA189, VEGFA206, and VEGFAx (7). It plays a central role in both physiological and pathological conditions, influencing tissue growth, repair, and the response to hypoxia. In addition to its vascular roles, VEGFA impacts several organs, including the kidney, lung, liver, and central nervous system (6).

VEGFA primarily binds to VEGFR2 on endothelial cells, initiating signaling pathways that promote cell survival, migration, and proliferation, all essential for blood vessel formation (7). Its regulation is tightly controlled, and dysregulation of VEGFA is implicated in diseases such as cancer, diabetic retinopathy, and age-related macular degeneration, where abnormal blood vessel growth contributes to disease progression. Upon binding to its receptor, VEGFA induces VEGFR dimerization, while specific tyrosine residues are phosphorylated in the intracellular region. This process recruits and activates intracellular signaling components. Consequently, several strategies for treating angiogenesis-dependent diseases target VEGFA signaling by blocking the interaction between VEGFA and VEGFR or inhibiting VEGF release (7,8).

The interaction between VEGFA and VEGFR has been resolved at the molecular level, including their binding mode (9), which is essential for the rational design of inhibitor molecules. Under physiological conditions, VEGFA exists as a homodimer, where two monomers are linked via disulfide bonds in an antiparallel arrangement (10,11). This structural configuration accommodates two receptor-binding sites. Co-crystal structures of the VEGFA-VEGFR complex reveal that VEGFA binds to two receptor molecules at the immunoglobulin-like domains D2, D3, and D2’, D3’, respectively (9). The specific residues involved in the interaction interface have also been identified. VEGFA antibodies have been developed to compete for these receptor-binding sites and inhibit VEGFA’s interaction with VEGFR. To date, numerous VEGF antibodies have been developed and used for this purpose (12–16).

Anti-VEGFA drugs include antibodies and soluble receptors, which are relatively large biomolecules. Widely used drugs in this category include bevacizumab, pegaptanib, ranibizumab, and aflibercept (12–16). Despite their effectiveness, these treatments have limitations. Therefore, peptide-based anti-VEGF drugs are under development. Peptides offer several advantages over both small molecules and large antibodies. Compared to small molecules, peptides are less toxic and degrade efficiently in the body, avoiding accumulation due to rapid protease activity and elimination. Advanced biomaterials and drug delivery systems enable a sustained release of peptides in the human body (17). Furthermore, their half-life can be extended through modifications such as the incorporation of non-natural amino acids, acylation, pegylation, cyclization, or the creation of peptidomimetics (7,18).

While antibodies have higher binding affinities, peptides can be administered at higher molar concentrations, mitigating issues related to high molecular weight, solubility, and aggregation tendencies (19,20). Additionally, unlike antibodies, which often require low-temperature storage, peptides provide greater formulation flexibility and stability, making them ideal for controlled drug delivery systems (7). However, these therapies still face challenges, as therapeutic outcomes are significantly influenced by the composition and degradation rate of the biomaterials used.

Various peptide-based VEGFA inhibitors have shown promise in modulating angiogenesis. Peptides that mimic glycosaminoglycans have demonstrated angiogenic effects, though their structural heterogeneity poses challenges for pharmaceutical development (21,22). In contrast, VEGFR-mimicking peptides bind effectively to VEGFA, with Je-7 inhibiting endothelial cell proliferation at an IC_50_ value of 0.1 µM (23). Further modifications to D,L-peptides have enhanced their stability and affinity, enabling them to block VEGFA interactions at nanomolar concentrations (24,25). Additionally, arginine-rich peptides inhibited VEGFA binding with an IC_50_ value of 2 µM and exhibited anti-angiogenic effects, reducing neovascularization and tumor metastasis (26). Stability enhancements, such as the use of all-D amino acid derivatives, maintained strong binding properties and extended half-life, while cyclic frameworks further improved anti-angiogenic potency. Peptides v107 and v114, with IC_50_ values of 0.70 µM and 0.22 µM respectively, effectively competed with VEGFA receptors for binding. Modifications to these peptides enhanced their binding affinity and enzymatic stability (7,27).

To discover more potent binders, a series of molecules was generated using the phage display method, building on earlier research to reengineer the Z-domain from staphylococcal protein A (28,29). The Z-domain comprises 58 amino acid residues arranged in three helices. In this approach, Z-1-2 was identified as the tightest-binding clone of the Z-domain peptides, featuring 59 residues with an additional alanine (30). Z-1-2 exhibited an IC_50_ of 0.34 µM to VEGFA. Further stabilizing mutations of the Z-domain led to an even smaller, 34 amino acid long peptide, with a two-helix structure and a stabilizing intramolecular disulfide bridge. From this scaffold, the mini-Z-1 peptide was derived, showing the highest inhibitory effect with a IC_50_ of 0.23 µM (30).

Recently, computational methods for peptide design, especially from the field of artificial intelligence (AI), have shown remarkable successes in developing peptide binders towards different target proteins (31–35). These methods could potentially provide much faster and cheaper routes to novel binders that target specific surface patches and have desired biophysical properties. Deep learning protein design methods, such as ProteinMPNN (36), can accurately predict sequences that have a high probability to fold into the desired target structure and/or bind to the desired receptor protein.

In this study, we took advantage of these new computational design tools in order to search for improved peptide-based VEGFA inhibitors. The mini-Z-1 peptide represents a good starting template for this goal, due to its promising affinity and anti-angiogenic activity, although its properties need to be further optimized. We aimed to achieve this goal by efficiently screening the vast sequence space with the help of computational design tools, while focusing the testing on the most promising peptide sequences.

## Results and Discussion

An overview of our combined computational and experimental pipeline is shown in Figure 1. Mini-Z peptides were developed in a stepwise approach as described below, leveraging Rosetta and ProteinMPNN tools for design as well as the Surface Plasmon Resonance (SPR) method for binding affinity testing.

**Figure 1:**
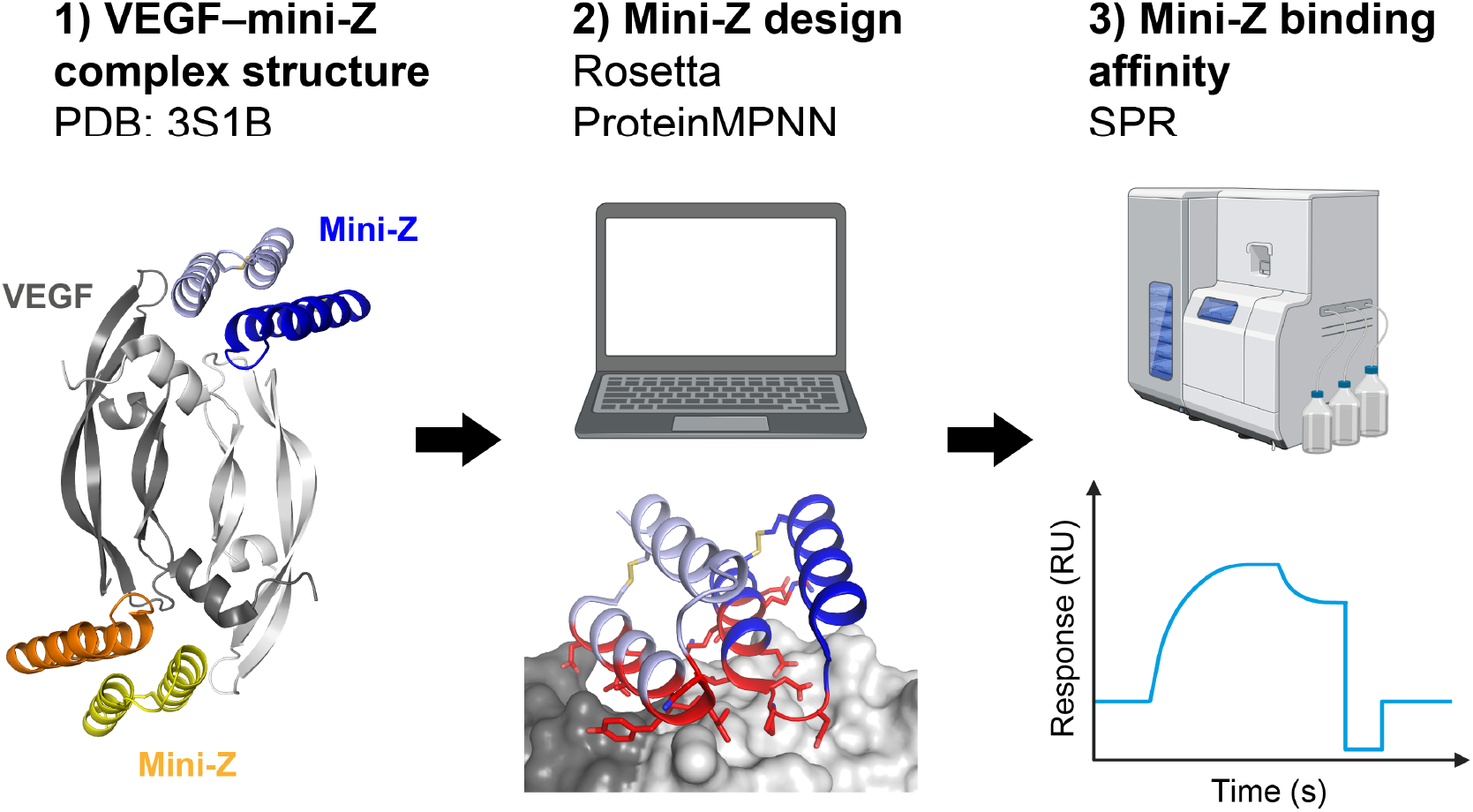
Workflow of designing mini-Z peptides. The 3D structure of the VEGF-mini-Z complex was obtained from PDB (step 1) and used as input to Rosetta and ProteinMPNN design calculations (step 2). Designed mini-Z peptide variants were afterwards tested for their binding affinity (step 3) using SPR.

### Characterization of the native mini-Z peptide

As the starting point of this study, we used the anti-VEGF peptide discovered previously by Fedorova et al. (30). This 34 amino acid long, so-called mini-Z peptide was originally derived from the Z-domain of staphylococcal protein A and optimized by several rounds of phage display (30). Critical for the stabilization of the peptide’s helix-turn-helix structure is an intra-molecular disulfide bond between Cys5 and Cys34.

We started with the characterization of the mini-Z peptide by calculating its Rosetta binding energy and measuring its binding affinity using SPR. A Rosetta interface (dG_separated) score of -106.77 Rosetta energy units (REU) and a K_D_ value of 9.27 ± 1.32 µM (Table 1 and Figure 2) indicated a favorable interaction of mini-Z with VEGF.

**Table 1.**
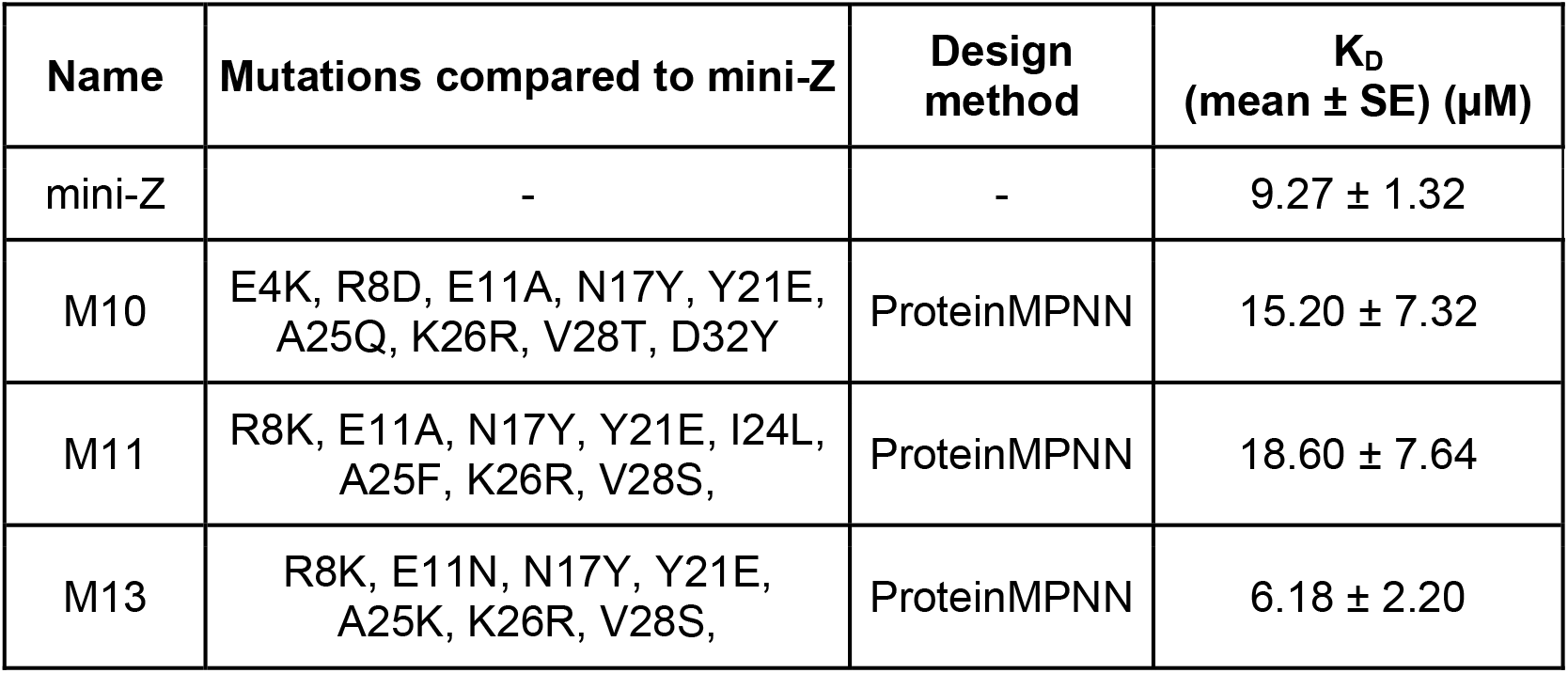
Summary of mutations and binding affinity constants of redesigned mini-Z peptides.

| Name | Mutations compared to mini-Z | Design method | $K_D$<br>(mean ± SE) (μM) |
| --- | --- | --- | --- |
| mini-Z | - | - | 9.27 ± 1.32 |
| M10 | E4K, R8D, E11A, N17Y, Y21E, A25Q, K26R, V28T, D32Y | ProteinMPNN | 15.20 ± 7.32 |
| M11 | R8K, E11A, N17Y, Y21E, I24L, A25F, K26R, V28S, | ProteinMPNN | 18.60 ± 7.64 |
| M13 | R8K, E11N, N17Y, Y21E, A25K, K26R, V28S, | ProteinMPNN | 6.18 ± 2.20 |

**Figure 2:**
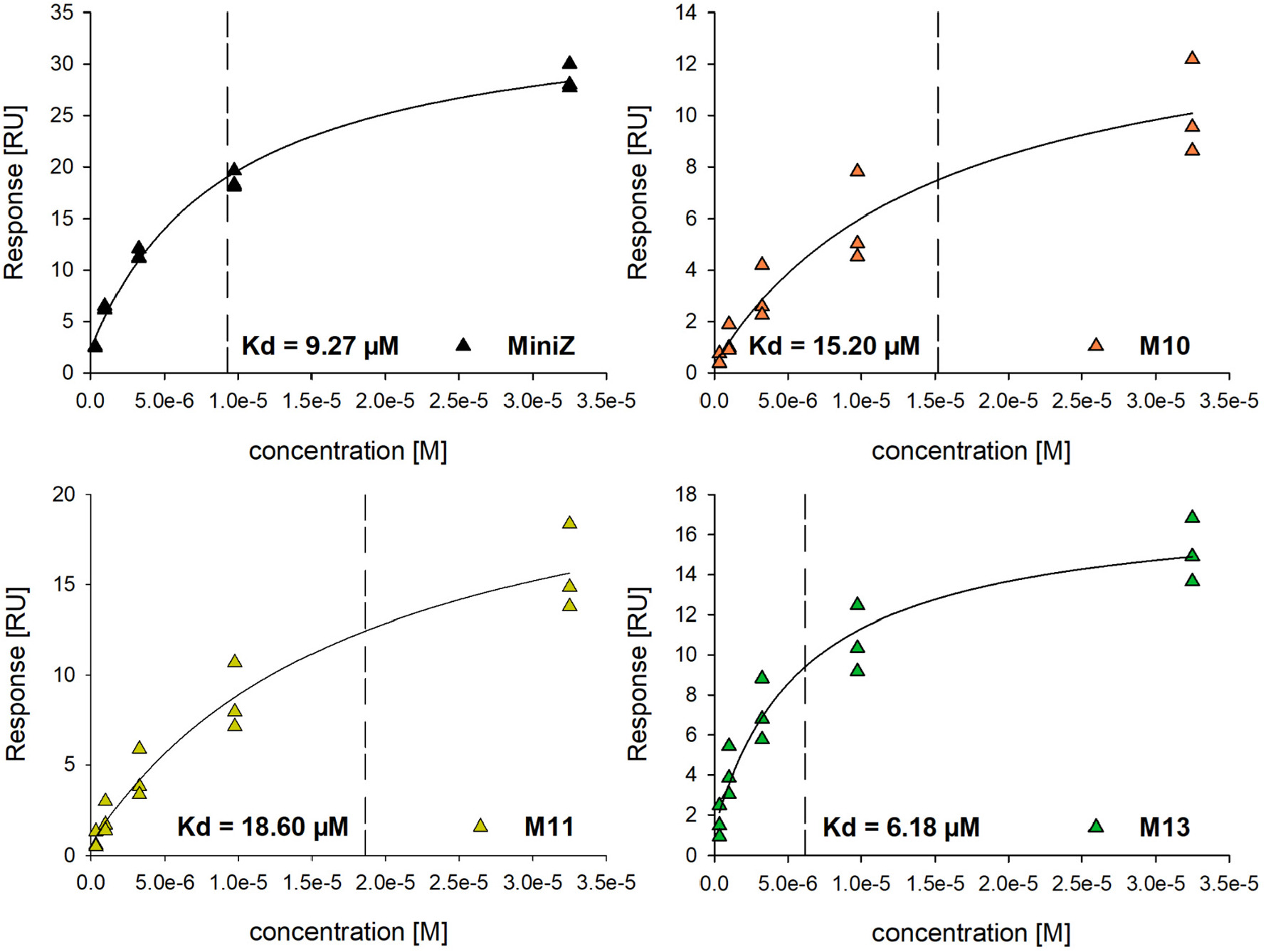
Affinity of original mini-Z and modified peptides M10, M11, M13 towards immobilized VEGF. Surface plasmon resonance measurements were performed for each peptide as a single cycle kinetic with increasing concentrations in triplicates. Here the steady state responses (R_eq_) were plotted against the respective peptide concentration (c). The K_D_ was determined by regression analysis using the steady state affinity model according to formula: R_eq_=(c□ R_max_)/(K_D_+c)+offset.

### Redesign of mini-Z using Rosetta and ProteinMPNN

To search for improved variants of the mini-Z peptide, which could offer a better binding affinity and inhibitory efficacy than the original peptide, we redesigned the VEGF-binding surface on mini-Z. We used a computational design strategy because this allowed screening several thousand peptide sequences, while focusing the experimental testing on the most promising candidates. A detailed description of the computational design workflow is provided in Methods and displayed in Supporting Figure S1.

The first set of design experiments used the Rosetta FixBB protocol, which generated 11 different peptide sequences. All of them showed equal or better in silico binding energy scores than mini-Z (see Supporting Table S1). Based on visual inspection of the 3D models of the designed mini-Z peptides bound to VEGF, we found two mutants most promising. The first one, mini-Z-M1, contains the mutations D15N and V28T. D15N introduces a new intra-molecular hydrogen bond to E11, thereby stabilizing the mini-Z structure, and V28T creates a new hydrogen bond to the amino group of K74 in VEGF (residue numbering according to isoform VEGF165 with Uniprot ID P15692). The second peptide, mini-Z-M2, carries the mutations R8L and D15N. The R8L mutation is present in almost all designs generated with the Rosetta FixBB protocol, as can be seen in the sequence logo in Figure S2A. The role of this mutation may be to stabilize the mini-Z structure via intra-molecular packing interactions with I30 as well as to reduce electrostatic repulsion with a nearby R131 in VEGF. In addition to the M1 and M2 mutants, two more peptides were designed based on the mutations sampled with Rosetta FixBB. The mini-Z-M3 peptide was obtained by combining the mutations from mini-Z-M1 and mini-Z-M2 (R9L, D15N, V28T), and the mini-Z-M4 peptide was designed by introducing the N17Y mutation into mini-Z-M1. This was done because calculations with the Rosetta FastDesign protocol suggested that the N17Y mutation could introduce an additional hydrogen bond interaction with Q48 in VEGF as well as van-der-Waals interactions with H53 in VEGF.

However, since the sequence space sampled with the Rosetta FixBB protocol was relatively small, we carried out a second set of design experiments using the ProteinMPNN tool, which generated 310 different sequences. To focus testing on the most promising peptides, we predicted their structures in complex with VEGF using AlphaFold3 (37) and filtered out all peptides which had low confidence scores (pLDDT-complex < 0.9 and pLDDT-mini-Z < 0.85) and a root-mean-squared deviation (RMSD) to the X-ray structure of VEGF-mini-Z of more than 0.6 Å. The remaining 168 peptides were visually inspected and all of them had better in silico binding energy scores than the original mini-Z (Table S1). In order to consider peptides with diverse sequences for testing, they were clustered into 15 groups based on their sequence differences, with a maximum of 3 amino acid differences between the sequences in order to form a cluster. Finally, we picked 10 peptides from different clusters (referred to as mini-Z-M5 to mini-Z-M14) to inspect them in closer detail. The 10 ProteinMPNN-designed peptides and the 4 Rosetta-designed peptides are considered good candidates for further follow-up testing and their sequences are summarized in Supporting Table S2.

Looking at the ProteinMPNN predicted amino acid probabilities (Figure S2B) and generated mini-Z sequences (Table S1), the following trends can be seen: In case of residues L7, K10, L14, P16, N19, and L20 only the original amino acid is predicted, whereas for the other interface residues their identity is changed. E4 can be mutated to lysine or arginine; R8 can be changed to leucine, aspartate or glutamate; E11 can be mutated to alanine or asparagine; Y21 can be replaced by glutamine or glutamate; I27 can be mutated to leucine; and D32 can be substituted to tyrosine or phenylalanine. The two most variable residues are A25, which can be mutated to either glutamate, glutamine, lysine or phenylalanine, and V28, which can be changed to either alanine, serine, isoleucine, threonine, or glutamate. Interestingly, residue N17 is always replaced by tyrosine in all peptides, indicating that this is the most favorable amino acid at this position.

### Binding affinity of designed mini-Z peptides

Next, we proceeded with testing some of the designed mini-Z peptides, which we obtained with ProteinMPNN in the previous step (Table S2). For this purpose, the linear mini-Z peptides M10, M11, and M13 were synthesized via solid-phase synthesis (see details in the Methods) and cyclized through the formation of the conserved disulfide bridge between Cys5 and Cys34. The resulting cyclic peptides were purified using high-performance liquid chromatography, achieving a purity of over 90%. The identity of the sequence and the completeness of disulfide bridge formation were confirmed via mass spectrometry and chemically using DTNB, respectively.

The analysis of binding kinetics and affinities was performed using SPR, wherein VEGF was immobilized, and the mini-Z, the designed mini-Z variants or Bevacizumab were examined as soluble ligands using a Single Cycle Kinetic approach with increasing concentrations. The mini-Z peptide exhibited strong binding to VEGF, with a K_D_ value of 9.27 ± 1.32 µM (Table 1 and Figure 2). Subsequent analysis of the peptide variants M10, M11, and M13 yielded comparable values of 15.20 ± 7.32 µM (M10) and 18.60 ± 7.64 µM (M11), alongside a significantly improved binding strength of 6.18 ± 2.20 µM (M13). A more detailed analysis of the SPR sensorgrams (Figure S3) indicated that the enhanced binding was attributed to reduced dissociation.

### Structural basis for enhanced VEGF interaction of mini-Z-M13

Comparing the mini-Z structure with the designed model of M13 (Figure 3A) provides some hints on the structural mechanisms that could contribute to the higher binding affinity of M13. The sequence of M13 differs in 7 positions from that of mini-Z, and contains two mutations, E11N and A25K, which are unique to M13, but not observed in M10 or M11. As can be seen in Figure 3B, the native mini-Z residue E11 makes two hydrogen bonds (to VEGF-Y51 and to K26), which are changed upon mutation to asparagine in M13. This releases residue R26 from its interaction with N11 and allows the formation of a new intermolecular hydrogen bond between mini-Z and VEGF, which could stabilize the binding interface (Figure 3B). New interactions are also observed in case of the other mutations. The small-to-large mutation N17Y leads to increased van-der-Waals interactions with the loop connecting the α1 helix and β1 strand in VEGF (Figure 3C). Furthermore, the double mutation Y21E and A25K creates an intramolecular salt bridge clamp on the surface of the second helix of the mini-Z peptide, which could increase its overall stability (Figure 3D). E21 is also posed to interact with the sidechain of VEGF-K42 via a salt bridge (VEGF residue numbering according to isoform VEGF165 with Uniprot ID P15692). In summary, this analysis identifies several new interactions between VEGF and M13, which can be responsible for its improved affinity.

**Figure 3:**
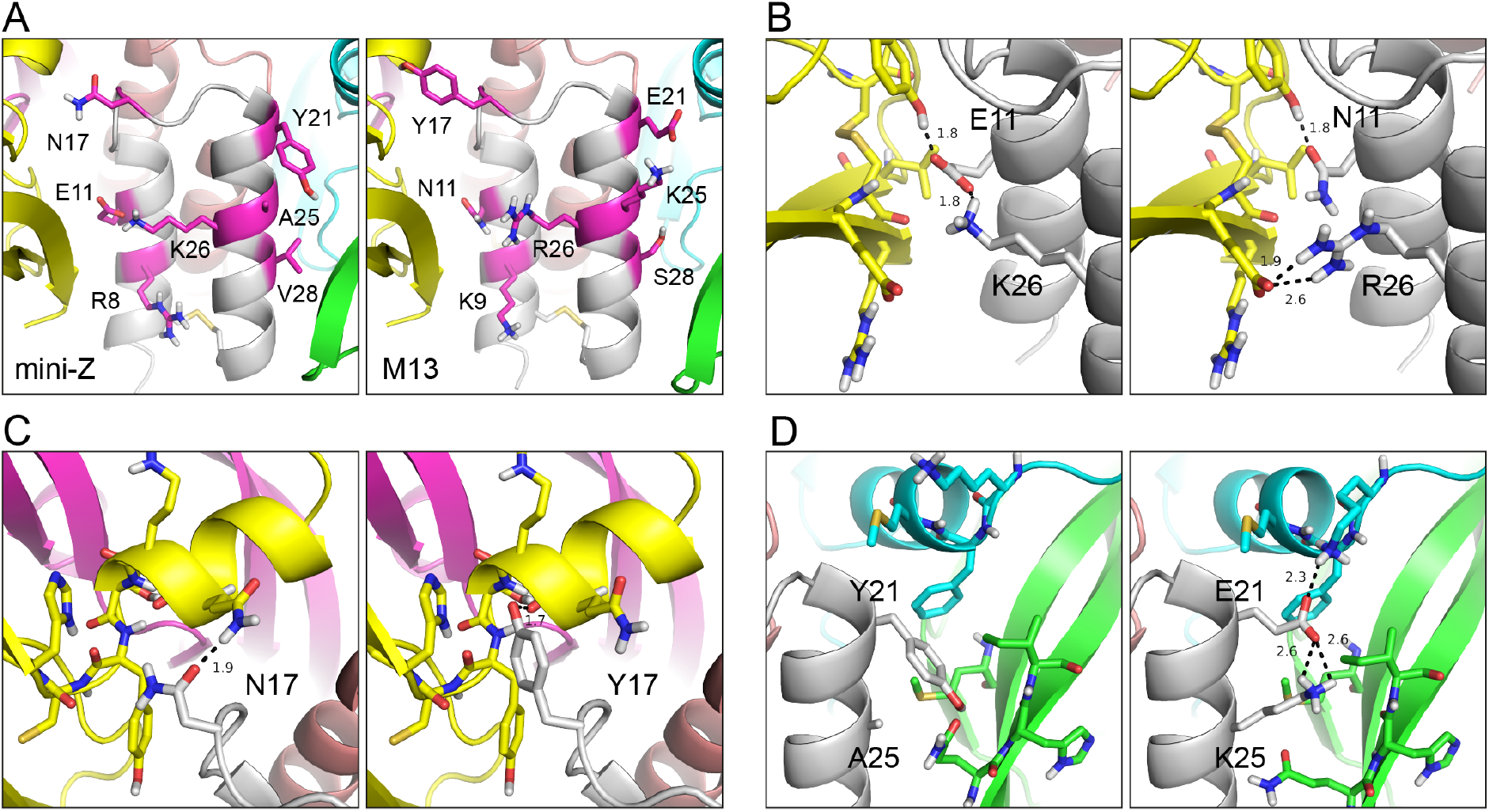
Structural models of mini-Z and its variant M13 showing designed mutations. (A) Residues in mini-Z (left) which were mutated in M13 (right) are shown as sticks and colored magenta. (B) Binding sites of E11 and K26 in mini-Z (left) and of the mutated residues N11 and R26 in M13. (C) Binding interface of N17 in mini-Z (left) and of the mutated residue Y17 in M13 (right). (D) Binding sites of Y21 and A25 in mini-Z (left) and of the mutated residues E21 and K25 in M13. Potential hydrogen bonds are indicated with dashed lines.

## Conclusion

In conclusion, we could demonstrate the redesign of the mini-Z peptide to enhance its binding affinity against VEGFA. With the help of Rosetta and ProteinMPNN tools, several promising variants were generated, with M13 showing a pronounced binding improvement, with a K_D_ of 6.2 µM compared to a K_D_ of 9.3 µM for the native mini-Z. The study also demonstrates that the used pipeline enables obtaining peptide binders in one round of design and testing.

## Materials and Methods

### Rosetta design calculations

The crystal structure of the dimeric receptor binding domain of VEGFA (corresponding to residues 218-313 of isoform VEGF189, Uniprot ID P15692) bound to mini-Z was downloaded from the Protein Data Bank under accession number 3S1B (30). In order to prepare the structure for the subsequent Rosetta design calculation, it was energy-minimized using the Rosetta ref2015 energy function (38), optimizing all side chain degrees of freedom while applying harmonic constraints to the protein backbone atoms. Because mini-Z binds as a symmetric homodimer to the VEGF dimer, a symmetry definition file was created to fix the internal symmetry of mini-Z during the Rosetta calculations. The symmetric system comprised one mini-Z dimer flanked on opposite sides by two VEGF dimers, a first one from the asymmetric unit and a second one representing a crystal symmetry mate. The central C2 symmetry axis of the system was set to the mini-Z dimer axis.

Design calculations on mini-Z were carried out using the Fixbb and FastDesign protocols available in Rosetta (version 3.14) (39). In total, 1000 design calculations, starting with different random seeds, were performed, yielding 36 unique sequences. The designs were sorted by the Rosetta binding energy score (dG_separated) and binding energy density (dG_separated / dSASA) for mini-Z and visually analyzed in the PyMOL program. Four mini-Z sequences were selected for experimental testing.

### ProteinMPNN design calculations and selection of designed peptides

An HPC compatible version of ProteinMPNN (36) (https://github.com/dauparas/ProteinMPNN) was used with the following settings: the ‘chains_to_design’ flag was set to the two chains comprising the mini-Z dimer and was used in combination with the ‘design_only_position’ flag to sample only the following mini-Z residues that are at the interface with VEGF: E4, L7, R8, K10, E11, L14, P16, N17, N19, L20, Y21, I24, A25, K26, V28, D31, D32. The ‘tied_positions’ flag was turned on to ensure that changes made to one mini-Z chain were mirrored to the second chain to maintain the symmetry of the mini-Z dimer. Finally, the sampling temperature was set to 0.1.

1000 ProteinMPNN calculations with different random seeds were performed, yielding 310 different mini-Z sequences. To select the most promising designs for experimental testing, the following procedure was applied: (1) AlphaFold3 (37) web server was used to predict the structure of the VEGF-mini-Z complex for every new mini-Z sequence. Those designs which yielded an AlphaFold model with an average pLDDT >= 90 over the entire complex and an average pLDDT >= 85 over all mini-Z residues were kept, yielding 168 acceptable sequences. We ensured that all 168 AlphaFold models had a low RMSD (of maximal 0.6 Å) to the VEGF-mini-Z crystal structure (30). (2) The mutations from each accepted sequence were introduced in the VEGF-mini-Z crystal structure using the Rosetta MutateResidueMover and the binding energy score between mini-Z and VEGF (dG_separated) was calculated for each designed protein. (3) Afterwards, sequence clustering with a sequence difference cutoff of 2 amino acids was applied to reduce redundancy in the designs, yielding 53 clusters. (3) From each cluster, the model with the best dG_separated score was selected and visually analyzed in the PyMOL program [The PyMOL Molecular Graphics System, Version 2.5.4, Schrödinger, LLC]. The structural models were checked for features like tightly packed side chains at the mini-Z-VEGF interface and favorable salt bridge and hydrogen bond interactions, and all designs were found acceptable at that step. (4) Finally, the 53 checked designs were clustered again using this time a sequence difference cutoff of 3 amino acids. From the obtained 15 clusters, a maximum of 1 designed sequence per cluster and a total of 10 mini-Z sequences were selected for experimental testing.

### Chemicals

Fmoc-and Boc-protected amino acids were obtained from Sigma Aldrich and used as purchased. Fmoc-protected cysteine-preloaded Wang resin (substitution level S = 0.333 mmol/g) was obtained from Sigma Aldrich. DIPEA (N,N-Diisopropylethylamine) and HATU (O-(7-Azabenzotriazol-1-yl)-N,N,N′,N′-tetramethyluronium-hexafluorphosphat) were obtained from BLD Pharm. Trifluoroacetic acid (TFA) and acetic anhydride were manufactured by Carl Roth. DMF was obtained from Iris Biotech, triisopropylsilane (TIS) from BLD Pharm, and 1,2-ethanedithiol (EDT) from Sigma Aldrich.

### Peptide synthesis

Peptides were synthesized using the solid-phase peptide synthesis (SPPS) strategy on an automated peptide synthesizer, Multipep 1. Initially, a preloaded Fmoc-protected cysteine Wang resin was deprotected using 20% 4-methylpiperidine in DMF (10 min x 2). The second amino acid (4 equivalents relative to the preloaded amino acid) was dissolved in DMF, preactivated for 5 min using HATU and DIPEA, and subsequently transferred onto the resin to react for 1.5 hours. These steps were repeated iteratively for the entire peptide sequence, with intermediate acetylation steps using acetic anhydride (5% in DMF) to cap delayed sequences. The resin-bound peptide was manually cleaved using a cleavage cocktail comprising 92.5% trifluoroacetic acid (TFA), 2.5% triisopropylsilane (TIS), 2.5% ethanedithiol (EDT), and 2.5% Milli-Q water. The linear peptides were precipitated from the cleavage cocktail using cold diethyl ether, centrifuged, and isolated by decantation. The crude peptide was washed multiple times with cold ether to remove residual cleavage cocktail components.

Linear sequences were cyclized via disulfide bond formation using 30% hydrogen peroxide in water. The crude peptide was dissolved or suspended in Milli-Q water (with a small amount of methanol added for highly lipophilic sequences). The pH was adjusted to ∼8 using ammonium hydroxide, and the reaction mixture was left to oxidize cysteine thiols to disulfide bridges over 30 minutes. The reaction was quenched by dropwise addition of TFA until the pH was adjusted to 3–4. The mixture was then frozen and lyophilized, yielding crude cyclized products. These crude cyclized peptides were purified using preparative high-performance liquid chromatography (HPLC) with a reverse-phase column, employing a gradient of 5% to 95% acetonitrile in water and afterwards dried using vacuum centrifugation.

The peptides were resolubilized in 10 mM TEAB (triethylammonium bicarbonate buffer; pH 8.5; Sigma-Aldrich; T7408) with the assistance of ultrasound. To remove residual contaminants from the synthesis that could interfere with subsequent analysis, a buffer exchange and peptide purification were performed using gel filtration chromatography (PD MiniTrap G-10 columns; Cytiva; 28918010). The buffer exchange was conducted to HBS-EP buffer pH 7.4 (Cytiva; BR-1008-26). The final concentration was determined by UV absorption measurement at 280 nm, taking into account the extinction coefficient and molecular weight of the respective peptide using a spectrophotometer (Nanodrop 2000, Thermo Fisher).

### Surface Plasmon Resonance measurements

The binding analysis was conducted through measurements by SPR using a Biacore T200 device (Cytiva). VEGF165 (Promega; J2371) was immobilized on a Series S Sensor Chip CM5 (Cytiva; 29104988) using an Amine Coupling Kit (Cytiva; BR100050) and 10 mM potassium acetate pH 5.5 + 0.05% Tween20 as the immobilization buffer. The immobilization level of VEGF at 542 RUs allows for maximum binding of the mini-Z peptides in the range of 50 RUs. For the measurement of interaction, concentrations ranging from 325 nM to 32.5 µM were prepared, encompassing five dilution steps of mini-Z and sequence-variant peptides, and analyzed in a Single Cycle Kinetic format at a flow rate of 30 µl/min. The analysis consisted of consecutive injections lasting 240 s, followed by a final dissociation time of 600 s and a regeneration injection for 90 s using 10 mM glycine pH 2.0. The resulting data were double-referenced by subtracting the nonspecific signal from a flow cell without VEGF immobilization as well as the blank signal caused by the running buffer HBS-EP. Data analysis was performed with Biacore T200 Evaluation Software 3.2. For the regression of the steady-state affinity, the RUs measured 4 seconds prior to the injection stop, with a window of 5 seconds for each concentration, were utilized. Following the fitting of the steady-state affinity model of these RUs in relation to the concentration, the K_D_ value was derived according to the formula: R_eq_ = c x R_max_ / (K_D_ + c) + offset

R_eq_ Response in equilibrium

R_max_ maximum binding capacity of the surface

K_D_ Equilibrium dissociation constant

c Analyte concentration

offset Response at zero analyte concentration.

## Acknowledgements

S.K., J.M., and G.K. acknowledge funding from the European Innovation Council (project 101130454 -ISOS). G.K is further supported by the Deutsche Forschungsgemeinschaft (DFG) (project 421152132, CRC1423, subproject C07 and project 448298270, TRR 386, subprojects A2 and B2), by the Sächsische Aufbaubank (project 3001070284 within program JTF InfraProNet 2021-2027), and by the Federal Ministry of Education and Research of Germany and by Sächsische Staatsministerium für Wissenschaft, Kultur und Tourismus in the programme Center of Excellence for AI-research “Center for Scalable Data Analytics and Artificial Intelligence Dresden/Leipzig”, project identification number: ScaDS.AI. J.M. is supported by a Humboldt Professorship of the Alexander von Humboldt Foundation. J.M. acknowledges funding by the Deutsche Forschungsgemeinschaft (DFG) through SFB1423 (421152132), SFB 1664 (514901783), TRR (514664767), and SPP 2363 (460865652). J.M. is supported by the Federal Ministry of Education and Research (BMBF) through the Center for Scalable Data Analytics and Artificial Intelligence (ScaDS.AI), through the German Network for Bioinformatics Infrastructure (de.NBI), and through the German Academic Exchange Service (DAAD) via the School of Embedded Composite AI (SECAI 15766814). Work in the Meiler laboratory is further supported through the National Institute of Health (NIH) through R01 HL122010, R01 DA046138, R01 AG068623, U01 AI150739, R01 CA227833, R01 LM013434, S10 OD016216, S10 OD020154, S10 OD032234. SK further acknowledges funding from German Federal Ministry of Education and Research (Bundesministerium für Bildung und Forschung, BMBF) (project REGAGforBone, grant: 01KC2304C and SyMBoD, grant 01ZX2210B and 01ZX1910B). C.L. is supported by DFG (TRR 386, subprojects Z2), by the Sächsische Aufbaubank (project 985100-070 within program JTF InfraProNet 2021-2027), and by the Federal Ministry of Education and Research of Germany in the programme “Zielgerichtete Entwicklung neuer Speziallipide und deren Formulierung”, project identification number 16PL301202. The authors thank the Leipzig University Computing Center and the NHR Center of TU Dresden (projects p_sdslnmrdata and p_peptide) for providing the computational resources.

