## Supplemental Information for "Computer-guided design of Z domain peptides with improved inhibition of VEGF"

### Supporting Figures and Tables

**Figure S1**: Detailed workflow schema of computational design pipeline for mini-Z peptides.

**Figure S2:** Sequence logo from Rosetta FixBB and ProteinMPNN design calculations.

**Figure S3:** Sensorgram with representative binding of mini-Z and variants M10, M11, M13 to VEGF from a single cycle kinetic measurement.

**Table S1:** Sequences and prediction scores of mini-Z peptide mutants designed with ProteinMPNN or Rosetta FixBB, respectively.

**Table S2:** Sequences of mini-Z peptide mutants designed with ProteinMPNN or Rosetta FixBB, which were selected for experimental testing.

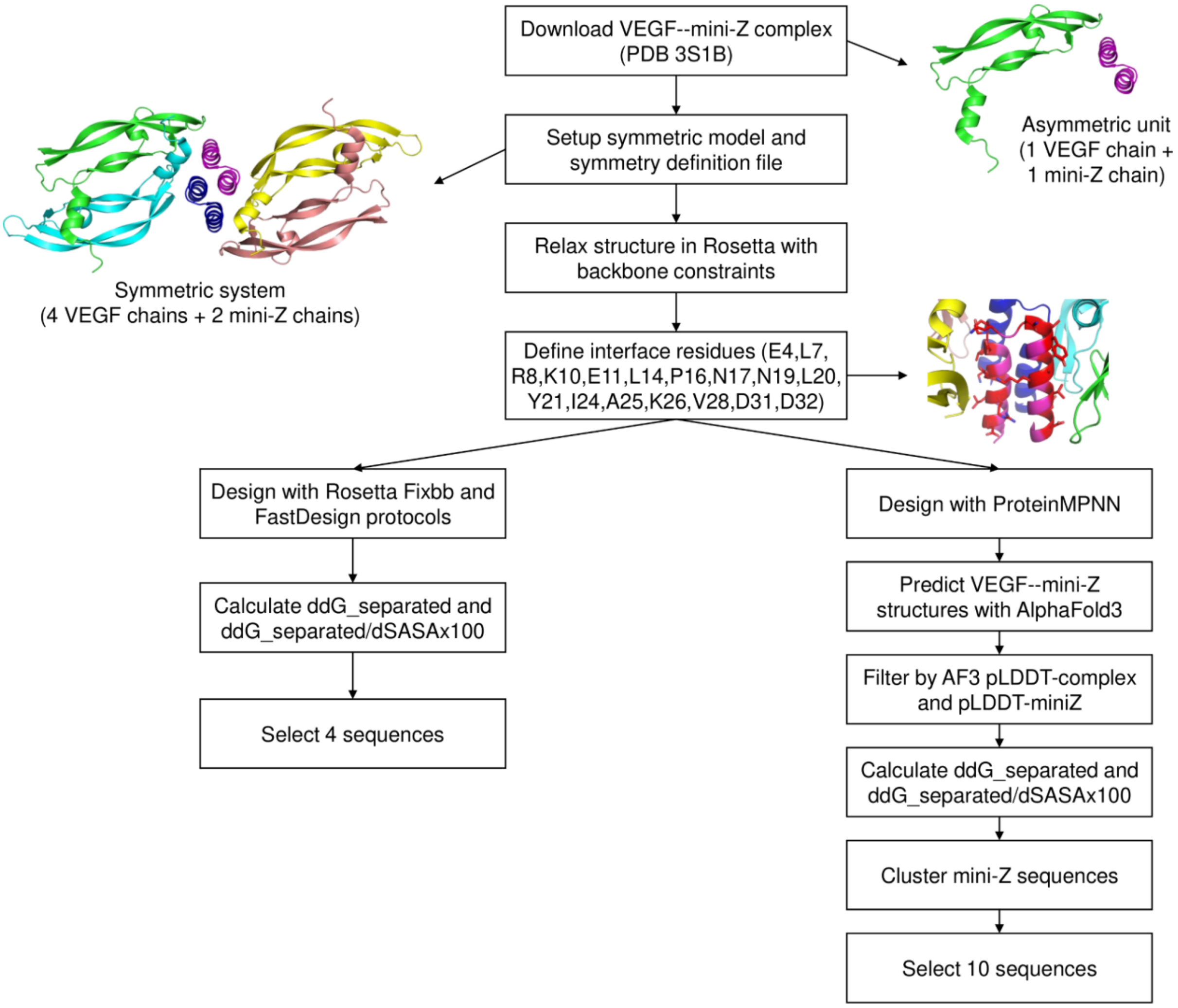


**Figure S1**: Detailed workflow schema of computational design pipeline for mini-Z peptides.

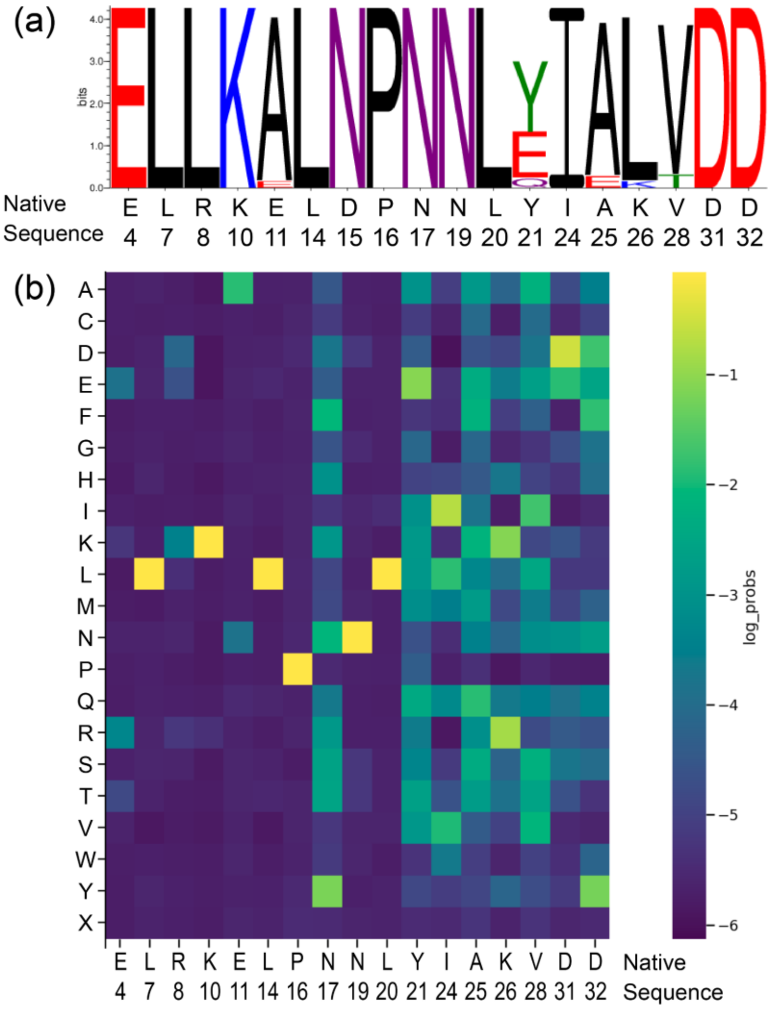


**Figure S2:** Sequence logo from Rosetta FixBB design calculations (a) and matrix of amino acid probabilities calculated by ProteinMPNN (b).

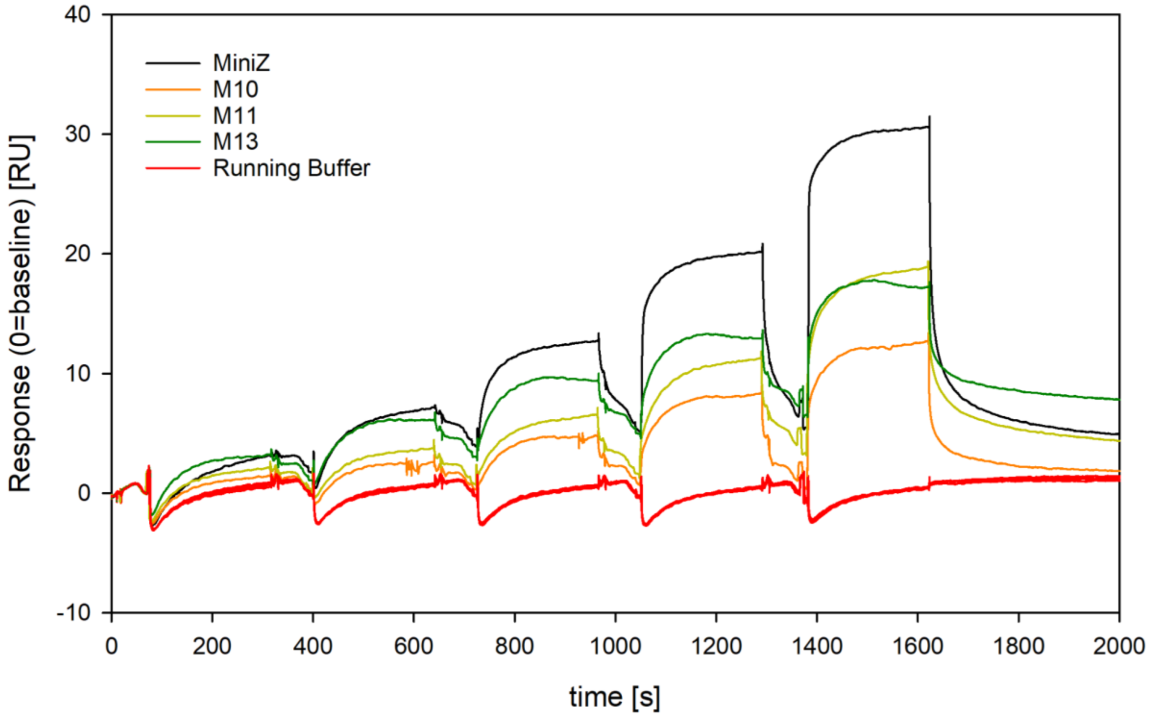


**Figure S3:** Sensorgram with representative binding of mini-Z and variants M10, M11, M13 to VEGF from a single cycle kinetic measurement. Shown is the binding in response units (RU) depending on increasing concentrations of the respective peptides. The background signal due to the running buffer is shown in red. The RUs at the end of each injection cycle (see marker) were used for $K_D$ determination by steady state affinity model.

**Table S1:** Sequences and prediction scores of mini-Z peptide mutants designed with ProteinMPNN or Rosetta FixBB, respectively. Sequences that were considered for testing are colored red and their scores are in bold.

| Name | Cluster-ID | pLDDT complex | pLDDT miniZ | RMSD to Xtal structure | dG_separated | dG_separated / dSASAx100 | Sequence |
|---|---|---|---|---|---|---|---|
| ProteinMPNN | | | | | | | |
| Native | | | | | -106.77 | -3.58 | FNKECLLRYKEAALDPNL NLYQRIAKIVSIDDDC |
| **3s1b_mpnn_96_0002** | **1** | **90.38** | **85.61** | **0.54** | **-131.44** | **-3.76** | FNKKCLLDYKNAALDPYL NLQQRIERIISIDYDC |
| 3s1b_mpnn_281_0001 | 1 | 91.00 | 86.48 | 0.56 | -128.84 | -3.70 | FNKRCLLDYKNAALDPYL NLQQRIERIASIDDDC |
| **3s1b_mpnn_99_0001** | **2** | **91.22** | **88.19** | **0.44** | **-130.15** | **-3.76** | FNKRCLLLYKAAALDPYL NLQQRIERIASIDDDC |
| 3s1b_mpnn_243_0001 | 2 | 90.30 | 86.44 | 0.52 | -134.73 | -3.79 | FNKRCLLLYKAAALDPYL NLQQRIERIISIDYDC |
| 3s1b_mpnn_139_0001 | 3 | 91.44 | 87.52 | 0.45 | -127.32 | -3.72 | FNKTCLLKYKAAALDPYL NLQQRLERIASIDDDC |
| 3s1b_mpnn_220_0001 | 3 | 90.46 | 86.75 | 0.43 | -121.29 | -3.78 | FNKECLLKYKAAALDPYL NLQQRIERIESIDDDC |
| 3s1b_mpnn_43_0001 | 4 | 90.15 | 85.10 | 0.49 | -120.35 | -3.76 | FNKECLLKYKNAALDPYL NLEQRIKKIISIDYDC |
| 3s1b_mpnn_198_0001 | 4 | 90.51 | 86.84 | 0.60 | -122.25 | -3.77 | FNKECLLRYKNAALDPYL NLEQRIKRIISIDYDC |
| **3s1b_mpnn_216_0001** | **4** | **90.29** | **84.85** | **0.54** | **-131.98** | **-3.70** | FNKRCLLEYKNAALDPYL NLEQRIKKIISIDYDC |
| 3s1b_mpnn_272_0001 | 4 | 90.50 | 86.39 | 0.59 | -133.76 | -3.74 | FNKRCLLLYKNAALDPYL NLEQRIKRIISIDYDC |
| 3s1b_mpnn_9_0001 | 5 | 90.13 | 85.09 | 0.55 | -129.38 | -3.72 | FNKKCLLDYKNAALDPYL NLEQRIQRIISIDYDC |
| **3s1b_mpnn_33_0002** | **5** | **90.75** | **85.96** | **0.52** | **-130.21** | **-3.63** | FNKRCLLDYKNAALDPYL NLEQRIQRIVSIDYDC |
| 3s1b_mpnn_79_0002 | 5 | 90.22 | 85.07 | 0.56 | -134.03 | -3.82 | FNKRCLLDYKNAALDPYL NLEQRIQRIISIDYDC |
| 3s1b_mpnn_105_0001 | 5 | 90.49 | 86.63 | 0.59 | -131.00 | -3.72 | FNKRCLLEYKNAALDPYL NLEQRIQRIISIDYDC |
| 3s1b_mpnn_169_0001 | 5 | 90.52 | 86.83 | 0.55 | -135.05 | -3.84 | FNKKCLLEYKNAALDPYL NLEQRIQRIISIDYDC |
| 3s1b_mpnn_130_0001 | 6 | 90.14 | 85.90 | 0.55 | -132.04 | -3.76 | FNKTCLLKYKNAALDPYL NLEQRIQRIASIDDDC |
| 3s1b_mpnn_285_0001 | 6 | 90.03 | 85.98 | 0.52 | -132.01 | -3.78 | FNKTCLLKYKNAALDPYL NLEQRIQRIISIDYDC |
| 3s1b_mpnn_135_0001 | 7 | 90.33 | 85.92 | 0.51 | -132.09 | -3.60 | FNKRCLLDYKAAALDPYL NLEQRIFRITSIDYDC |
| 3s1b_mpnn_233_0001 | 7 | 91.00 | 86.96 | 0.59 | -133.97 | -3.73 | FNKRCLLDYKNAALDPYL NLEQRIFRIESIDYDC |
| **3s1b_mpnn_29_0001** | **8** | **91.48** | **87.68** | **0.49** | **-126.16** | **-3.64** | FNKRCLLEYKAAALDPYL NLEQRIFRIVSIDYDC |
| 3s1b_mpnn_119_0001 | 8 | 91.80 | 88.70 | 0.42 | -129.33 | -3.69 | FNKRCLLEYKAAALDPYL NLEQRIKRIISIDDDC |
| 3s1b_mpnn_244_0002 | 8 | 91.54 | 87.94 | 0.40 | -131.27 | -3.77 | FNKRCLLEYKAAALDPYL NLEQRIKRIVSIDYDC |
| 3s1b_mpnn_261_0001 | 8 | 91.77 | 88.66 | 0.37 | -129.24 | -3.82 | FNKRCLLEYKAAALDPYL NLEQRIQRIVSIDYDC |

| | | | | | | | |
|---|---|---|---|---|---|---|---|
| 3s1b_mpnn_85_0001 | 9 | 90.83 | 86.66 | 0.46 | -130.03 | -3.77 | FNKKCLLDYKAAALDPYLNLEQRIQKIISIDYDC |
| **3s1b_mpnn_91_0001** | **9** | **91.12** | **87.27** | **0.49** | **-128.92** | **-3.68** | FNKKCLLDYKAAALDPYLNLEQRIQRIESIDYDC |
| 3s1b_mpnn_191_0001 | 9 | 91.21 | 87.69 | 0.43 | -127.06 | -3.62 | FNKKCLLDYKAAALDPYLNLEQRIKRIISIDYDC |
| 3s1b_mpnn_55_0002 | 10 | 91.23 | 88.18 | 0.41 | -118.75 | -3.68 | FNKECLLRYKAAALDPYLNLEQRIFRIISIDYDC |
| 3s1b_mpnn_57_0001 | 10 | 90.32 | 86.39 | 0.43 | -121.46 | -3.77 | FNKECLLKYKAAALDPYLNLEQRIKRIISIDYDC |
| 3s1b_mpnn_92_0001 | 10 | 90.80 | 86.81 | 0.52 | -117.63 | -3.59 | FNKECLLKYKAAALDPYLNLEQRIERISSIDYDC |
| 3s1b_mpnn_97_0001 | 10 | 90.72 | 86.41 | 0.41 | -123.36 | -3.78 | FNKECLLKYKAAALDPYLNLEQRIQKIISIDYDC |
| 3s1b_mpnn_118_0002 | 10 | 90.91 | 86.82 | 0.47 | -129.52 | -3.65 | FNKECLLKYKAAALDPYLNLEQRIQRIISIDYDC |
| 3s1b_mpnn_151_0002 | 10 | 90.44 | 86.14 | 0.41 | -117.28 | -3.74 | FNKECLLKYKAAALDPYLNLEQRIKRIISIDYDC |
| 3s1b_mpnn_156_0002 | 10 | 90.51 | 86.46 | 0.42 | -115.31 | -3.63 | FNKECLLKYKAAALDPYLNLEQRIFRITSIDYDC |
| **3s1b_mpnn_187_0002** | **10** | **90.74** | **86.29** | **0.42** | **-117.21** | **-3.72** | FNKECLLKYKAAALDPYLNLEQRLFRISSIDDDC |
| 3s1b_mpnn_190_0002 | 10 | 91.02 | 87.71 | 0.40 | -121.20 | -3.73 | FNKECLLKYKAAALDPYLNLEQRIQRIISIDYDC |
| 3s1b_mpnn_264_0001 | 11 | 90.90 | 87.18 | 0.35 | -135.65 | -3.89 | FNKTCLLKYKAAALDPYLNLEQRIKKIISIDYDC |
| 3s1b_mpnn_58_0001 | 12 | 91.07 | 87.72 | 0.60 | -123.01 | -3.49 | FNKNCLLKYKAAALDPYLNLEQRIKRIESINDDC |
| **3s1b_mpnn_168_0001** | **12** | **90.67** | **86.80** | **0.47** | **-114.85** | **-3.68** | FNKECLLRYKAAALDPYLNLEQRIKRIESIDDDC |
| 3s1b_mpnn_250_0001 | 12 | 90.27 | 85.87 | 0.51 | -128.89 | -3.65 | FNKTCLLKYKAAALDPYLNLEQRIKRIESIDDDC |
| **3s1b_mpnn_186_0001** | **13** | **90.15** | **84.55** | **0.51** | **-120.27** | **-3.66** | FNKECLLKYKNAALDPYLNLEQRIKRISSIDDDC |
| 3s1b_mpnn_267_0002 | 13 | 90.78 | 86.93 | 0.62 | -123.89 | -3.75 | FNKECLLKYKNAALDPYLNLEQRIFKIESIDDDC |
| 3s1b_mpnn_36_0001 | 14 | 92.09 | 89.07 | 0.43 | -130.96 | -3.74 | FNKRCLLLYKQAALDPYLNLEQRLQRIASIDDDC |
| 3s1b_mpnn_62_0001 | 14 | 90.39 | 86.00 | 0.48 | -129.50 | -3.78 | FNKRCLLDYKAAALDPYLNLEQRIQRIASIDYDC |
| 3s1b_mpnn_128_0001 | 14 | 90.62 | 86.94 | 0.49 | -118.15 | -3.73 | FNKECLLRYKAAALDPYLNLEQRIQRIASIDDDC |
| 3s1b_mpnn_133_0001 | 14 | 90.74 | 86.29 | 0.45 | -117.03 | -3.68 | FNKECLLKYKAAALDPYLNLEQRLQRIASIDDDC |
| 3s1b_mpnn_194_0002 | 14 | 91.92 | 88.61 | 0.45 | -126.31 | -3.76 | FNKRCLLDYKAAALDPYLNLEQRLERIASIDDDC |
| 3s1b_mpnn_230_0001 | 14 | 90.70 | 85.40 | 0.42 | -127.79 | -3.79 | FNKRCLLDYKAAALDPYLNLEQRIQRIASIDDDC |
| **3s1b_mpnn_252_0001** | **14** | **91.04** | **86.85** | **0.38** | **-125.03** | **-3.69** | FNKRCLLEYKAAALDPYLNLEQRLQRIASIDDDC |
| 3s1b_mpnn_306_0001 | 14 | 91.20 | 87.72 | 0.54 | -119.48 | -3.76 | FNKECLLKYKAAALDPYLNLEQRIQKIASIDDDC |
| 3s1b_mpnn_8_0002 | 15 | 90.41 | 85.50 | 0.59 | -127.52 | -3.63 | FNKKCLLDYKNAALDPYLNLEQRIFRISSIDDDC |
| 3s1b_mpnn_42_0002 | 15 | 91.38 | 87.22 | 0.35 | -126.86 | -3.67 | FNKRCLLEYKAAALDPYLNLEQRLFRISSIDDDC |
| 3s1b_mpnn_109_0001 | 15 | 90.83 | 86.62 | 0.58 | -130.82 | -3.77 | FNKRCLLDYKNAALDPYLNLEQRIFKISSIDDDC |

| | | | | | | | |
|---|---|---|---|---|---|---|---|
| 3s1b_mpnn_138_0001 | 15 | 91.35 | 87.75 | 0.58 | -128.03 | -3.62 | FNKRCLLEYKNAALDPYL NLEQRIFRISSIDDDC |
| FixBB | | | | | | | |
| Native | | | | | -99.95 | -3.53 | FNKECLLRYKEAALDPNL NLYQRIAKIVSIDDDC |
| 3s1b_fixbb_1 | | | | | -99.38 | -3.56 | FNKECLLLYKAAALNPNL NLYQRIALIVSIDDDC |
| 3s1b_fixbb_2 | | | | | -99.38 | -3.56 | FNKECLLLYKAAALNPNL NLEQRIALIVSIDDDC |
| **3s1b_fixbb_3** | | | | | **-102.15** | **-3.52** | FNKECLLLYKEAALNPNL NLYQRIAKIVSIDDDC |
| 3s1b_fixbb_4 | | | | | -102.15 | -3.52 | FNKECLLLYKEAALNPNL NLEQRIAKIVSIDDDC |
| 3s1b_fixbb_5 | | | | | -99.38 | -3.56 | FNKECLLLYKAAALNPNL NLQQRIELIVSIDDDC |
| 3s1b_fixbb_6 | | | | | -102.15 | -3.52 | FNKECLLLYKEAALNPNL NLQQRIEKIVSIDDDC |
| 3s1b_fixbb_7 | | | | | -99.38 | -3.56 | FNKECLLLYKAAALNPNL NLYQRIALIVSIDDDC |
| 3s1b_fixbb_8 | | | | | -98.70 | -3.53 | FNKECLLLYKAAALNPNL NLEQRIALIVSIDDDC |
| 3s1b_fixbb_9 | | | | | -98.96 | -3.54 | FNKECLLLYKAAALNPNL NLYQRIALITSIDDDC |
| 3s1b_fixbb_10 | | | | | -98.96 | -3.54 | FNKECLLLYKAAALNPNL NLEQRIALITSIDDDC |
| **3s1b_fixbb_11** | | | | | **-101.72** | **-3.50** | FNKECLLLYKEAALNPNL NLYQRIAKITSIDDDC |
| 3s1b_fixbb_12 | | | | | -101.72 | -3.50 | FNKECLLLYKEAALNPNL NLEQRIAKITSIDDDC |
| 3s1b_fixbb_13 | | | | | -98.94 | -3.54 | FNKECLLLYKAAALNPNL NLQQRIELITSIDDDC |
| **3s1b_fixbb_11 - R8L** | | | | | | | FNKECLLRYKEAALNPNL NLYQRIAKITSIDDDC |
| **3s1b_fixbb_11 - R8L +N17Y** | | | | | | | FNKECLLRYKEAALNPYL NLYQRIAKITSIDDDC |

**Table S2:** Sequences of mini-Z peptide mutants designed with ProteinMPNN or Rosetta FixBB, which were selected for experimental testing.

| Name | Design method | Mutations compared to mini-Z | Peptide Sequence | pI | Netcharge at pH 7.0 | grand average of hydropathicity |
|---|---|---|---|---|---|---|
| mini-Z | | - | FNKECLLRYKEAA LDPNLNLYQRIAKI VSIDDDC | 4.93 | -1 | -0.318 |
| M1 | Rosetta FixBB | D15N, V28T | FNKECLLRYKEA ALNPNLNLYQRIA KITSIDDDC | 6.23 | 0 | -0.462 |
| M2 | | R8L, D15N | FNKECLLLYKEAA LNPNLNLYQRIAK IVSIDDDC | 4.86 | -1 | -0.074 |
| M3 | | R8L, D15N, V28T | FNKECLLLYKEAA LNPNLNLYQRIAK ITSIDDDC | 4.86 | -1 | -0.218 |
| M4 | | D15N, N17Y, V28T | FNKECLLRYKEA ALNPYLNLYQRIA KITSIDDDC | 6.23 | 0 | -0.397 |
| M5 | Protein MPNN | E4K, R8D, E11N, N17Y, Y21Q, A25E, K26R, V28I, D32Y | FNKKCLLDYKNA ALDPYLNLQQRIE RIISIDYDC | 6.18 | 0 | -0.400 |
| M6 | | E4R, R8L, E11A, N17Y, Y21Q, A25E, K26R, V28A, | FNKRCLLLYKAAA LDPYLNLQQRIER IASIDDDC | 6.18 | 0 | -0.191 |
| M7 | | E4R, R8E, E11N, N17Y, Y21E, A25K, V28I, D32Y | FNKRCLLEYKNA ALDPYLNLEQRIK KIISIDYDC | 7.99 | +1 | -0.412 |
| M8 | | E4R, R8D, E11N, N17Y, Y21E, A25Q, K26R, D32Y | FNKRCLLDYKNA ALDPYLNLEQRIQ RIVSIDYDC | 6.18 | 0 | -0.426 |
| M9 | | E4R, R8E, E11A, N17Y, Y21E, A25F, K26R, D32Y | FNKRCLLEYKAA ALDPYLNLEQRIF RIVSIDYDC | 6.23 | 0 | -0.085 |
| M10 | | E4K, R8D, E11A, N17Y, Y21E, A25Q, K26R, V28T, D32Y | FNKKCLLDYKAA ALDPYLNLEQRIQ RITSIDYDC | 6.18 | 0 | -0.397 |
| M11 | | R8K, E11A, N17Y, Y21E, I24L, A25F, K26R, V28S | FNKECLLKYKAA ALDPYLNLEQRLF RISSIDDDC | 4.93 | -1 | -0.300 |
| M12 | | E11A, N17Y, Y21E, A25K, K26R, V28E | FNKECLLRYKAA ALDPYLNLEQRIK RIESIDDDC | 5.05 | -1 | -0.574 |
| M13 | | R8K, E11N, N17Y, Y21E, A25K, K26R, V28S | FNKECLLKYKNA ALDPYLNLEQRIK RISSIDDDC | 6.25 | 0 | -0.632 |
| M14 | | E4R, R8E, E11A, N17Y, Y21E, I24L, A25Q, K26R, V28A | FNKRCLLEYKAA ALDPYLNLEQRL QRIASIDDDC | 4.93 | -1 | -0.426 |
